# Reversing the Thyroid Hormone Mediated Repression of a HSV-1 Promoter through Computationally Guided Site Directed Mutagenesis

**DOI:** 10.1101/119115

**Authors:** Robert W Figliozzi, Feng Chen, Shaochung V Hsia

**Affiliations:** Department of Pharmaceutical Sciences, School of Pharmacy and Health Professions, University of Maryland Eastern Shore; Department of Natural Sciences, School of Agriculture and Natural Sciences, University of Maryland Eastern Shore

**Keywords:** Herpesvirus, Thyroid Hormone, Differentiation, Transcription Factor, Histone Modification

## Abstract

Thyroid hormones (TH or T_3_) and their DNA binding nuclear receptors (TRs), direct transcriptional regulation in different ways depending on the host cell environment and specific promoter characteristics of TH sensitive genes. This study sought to elucidate the impact on repression of nucleotide sequence/orientation of TR binding sites, TR elements, (TREs) within TH sensitive promoters. Computational analysis of the HSV-1 thymidine kinase (TK) gene TRE bound by TR and RXR revealed a single TRE point mutation sufficient to reverse the TRE orientation. In vitro experiments corroborated that the TRE point mutation exhibited distinct impacts on promoter activity, sufficient to reverse the TH dependent negative regulation in neuro-endocrine differentiated cells. EMSA and ChIP experiments suggest that this point mutation altered the promoter’s regulatory mechanism through discrete changes in transcription factor Sp1 and TR occupancy and altered enrichment of repressive chromatin, histone-3-lysine-9-trimethyl (H3K9Me3). Incites relating to this negative TRE (nTRE) mechanism impacts the understanding of other nTREs and TRE mutations associated with TH and herpes diseases.

## Introduction

Thyroid hormones (TH or T_3_) and their nuclear receptor family members Thyroid Hormone Receptors (TRs) have the ability to increase and decrease the rate of transcription of target genes (Lazar, 1993). These target genes generally contain a thyroid response element, TRE, arrangements of single or multiple DNA hexamers recognized and bound by the DNA binding domain (DBD) of TR (Yen, 2001). A host of criteria such as T_3_ binding to TRs, TR isoforms binding to target gene promoter regions as monomers, homo or hetero dimers, and the number of, arrangement and sequence of TREs determine the type of regulation (Yen, 2001). These criteria and their outcomes are extensively studied but still poorly understood for most thyroid hormone sensitive genes, which T_3_ decreased the target gene expression.

It is generally understood that certain TRE arrangements known as direct repeat fours, DR4, consensus AGGT(C/G)A nnnn AGGT(C/G)A, are bound by a TR and retinoid X receptor (RXR) heterodimer which recruit corepressor complexes to the target genes promoter region which modify the bound histones to repress transcription. When T_3_ as a ligand binds to the receptor a conformational change causes the corepressors to leave and be replaced by coactivators that modify the histones to attract the transcription machinery. Alternatively, several TREs with arrangements known as palindromes, often found on genes related to the feedback inhibition of T_3_ synthesis, regulate transcription in an opposite manner (Chatterjee et al., 1989). For these negative TREs (nTRE) the un-ligand TR somehow activated gene expression and then conferred repression upon binding T_3_ however this type of regulation is not well described possibly due to the conflicting circumstances for different genes that contain similar nTREs. There is argument over whether TR/DNA binding is maintained and what cofactors are involved because there has been evidence to support many different hypotheses depending on the system/cell line and gene being studied.

Our previous studies suggested that T_3_ participated in the HSV-1 regulation by repressing viral replication and gene expression (Bedadala et al., 2010;Figliozzi et al., 2014;Hsia et al., 2010). The HSV-1 Thymidine Kinase, TK, has been shown to play an intriguing role during reactivation (Kosz-Vnenchak et al., 1993;Nichol et al., 1996;Tal-Singer et al., 1997;Valyi-Nagy et al., 1994;Wilcox et al., 1992) and is one of the thyroid hormone sensitive genes that contains a palindromic nTRE which has seen some debate and intrigue for several decades (Maia et al., 1996;Park et al., 1993). The TK nTRE is in a context of palindrome with 6 nucleotides spacing (Pal6) (Hsia et al., 2010). In undifferentiated non-neuronal cells there is no regulation (Figliozzi et al., 2014;Hsia et al., 2010). However, it appears to result in TK being transcriptionally repressed by T_3_ in neuronal cells, evidenced by HSV-1 infection and in transfection reporter assay experiments (Hsia et al., 2010).

We have set out to better understand the precise TK nTRE nucleotide and TR/RXR residue interactions that define this protein-DNA binding, using web-based molecular biology applications to steer our benchtop TK nTRE site directed mutagenesis experiments. Performing a point mutation on a computationally identified nucleotide within the TRE, we generated a mutant that exhibited reversal of T_3_ sensitivity measured by dual luciferase assays. The luciferase reporter system has been used by several labs to evaluate nTREs in a variety of promoters including TSHα, TSHβ and HSV-1 TK (Lalli and Sassone-Corsi, 1995;Latif et al., 2016;Shibusawa et al., 2003). Electro-mobility shift assays (EMSA) were used to determine whether TR may exhibit differential binding to the wild type and mutant promoters. Chromatin Immuno-Precipitation (ChIP) Assays were used to determine what effects on the histone modification and SP-1 transcription factor binding that the mutations caused.

## Experimental Procedures

### Computational analyses of protein-DNA binding

The protein data bank (pdb) file 2NLL which depicts the crystallography structure of the RXR and TR heterodimer bound to a traditional TRE sequence 5'-CAG GTC ATT TCA GGT CAG-3' was manipulated using the Swiss PDB Viewer, SPV, (http://spdbv.vital-it.ch/) (Johansson et al., 2012) to obtain 2 separate pdb files for each protein monomer bound to a 5'-AGGTCA-3' hexamer TRE half-site. The interaction of TREs and nuclear receptors was computationally analyzed by the web based application PiDNA (http://dna.bime.ntu.edu.tw/pidn) (Lin and Chen, 2013). Each monomer-hexamer pdb was used as a template in the PiDNA web application to computationally predict the binding affinities of the monomers to the wild type hexamers and randomly generated single base pair mutant hexamers. A variety of criteria such as energy release index and position frequency matrix analyses. Hydrogen bond interactions between the DNA binding domain residues and hexamer nucleotides were measured using Python Molecule Viewer software (http://mgltools.scripps.edu/documentation/tutorial/python-molecular-viewer), Swiss PDB viewer, and PDBSum (http://www.ebi.ac.uk/thornton-srv/databases/cgi-bin/pdbsum/GetPage.pl?pdbcode=index.html) (Johansson et al., 2012). PDBSum calculates all hydrogen bonds between the DNA and the Protein in the pdb files. Hydrogen bonds were predicted using the HBPLUS (http://www.ebi.ac.uk/thornton-srv/software/HBPLUS/) default parameters were satisfied. The default parameters H-A distance is < 2.7, D-A distance is < 3.35, and the D-H-A angle is > 90. Schematic diagrams of protein-nucleic acid interactions were created by NUCPLOT (http://www.ebi.ac.uk/thornton-srv/software/NUCPLOT/) program (Luscombe et al., 1997;Luscombe et al., 1998).

### Construction of TRE mutant plasmid

The wild-type TK reporter plasmid pGL4.74-hRluc/TK was purchased from Promega (Cat # E6921). The single nucleotide mutation was introduced into the promoter region using Phusion Site-Directed Mutagenesis Kit from Thermo Fisher Scientific (Cat#: F-541) as described by the manufacturer. In short, template plasmid pGL4.74 was used in a PCR reaction using mutagenesis primers with 5’ phosphorylation modifications. The PCR reaction resulted in amplified linear point mutated plasmid in its entirety followed by a ligation to cyclize the linear PCR product to form circular plasmid containing single point mutation. The sequence and size of the generated plasmid were confirmed by gel electrophoresis and sequencing. The computational results discussed in a later section and depicted in Fig. 1 led to the development of the forward, 5’-tcgcatattaaggtgccgcgtgtggc-3’, and reverse, 5’-agtggacctgggaccgcgcc-3’, primer used for mutagenesis and strategy depicted in Fig. 2.

**Fig. 1:**
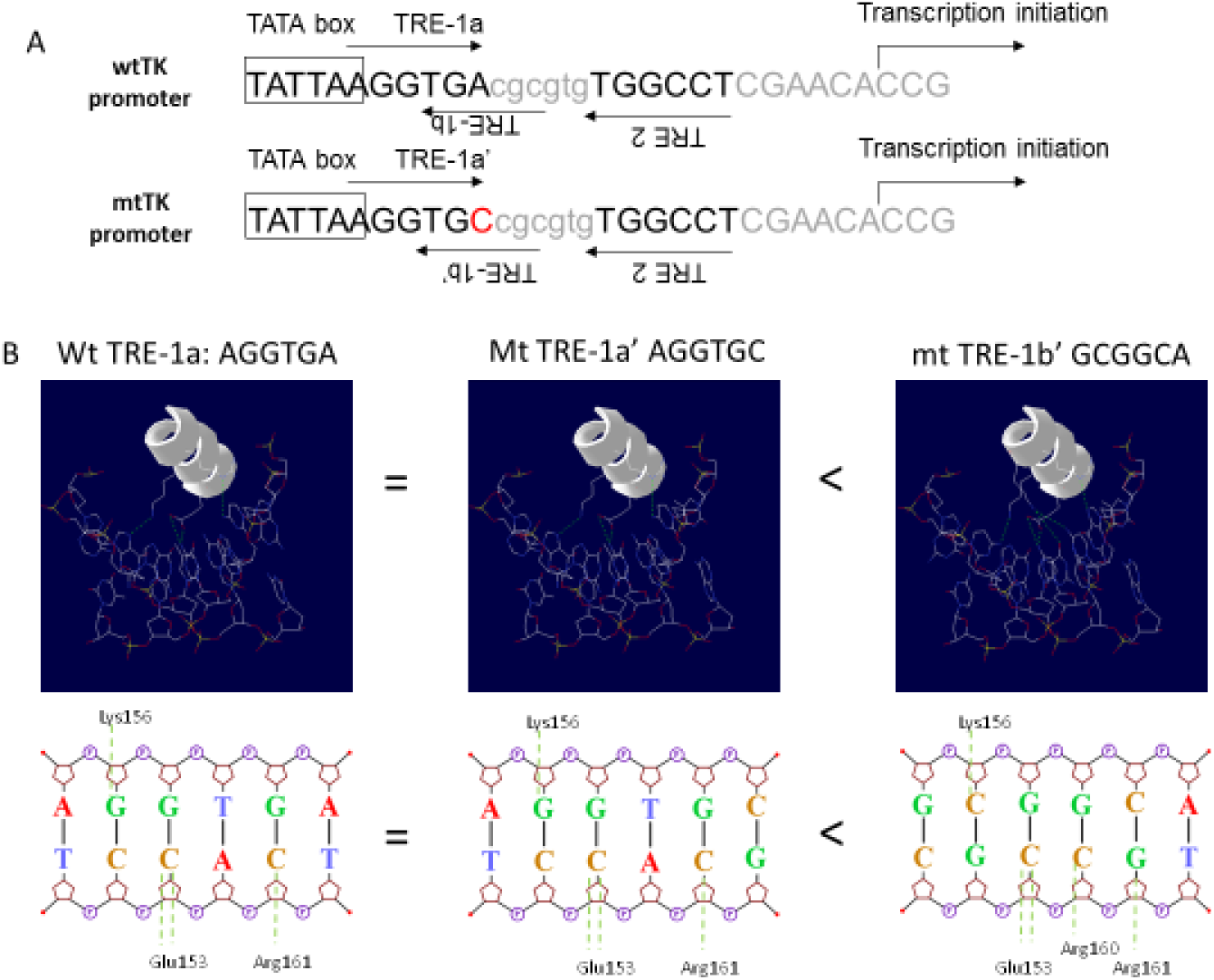
Computational analysis to predict binding of nuclear receptor to putative TK TREs. The interactions of protein and DNA were investigated by PiDNA, a web based application, to interpret the results from the biochemical assays. The high-scored putative TRE half-sites were further assessed for hydrogen bond interactions using Python Molecule Viewer (PMV) software and PDBSum. PMV was used to visualize and measure distances between the DNA binding protein residues and the bases of the DNA TRE half-sites. PDBSum calculates and diagrams putative hydrogen bonds between the DNA and the Protein in the pdb files. A. Random screening of PiDNA identified 5’-AGGTGA-3’ (TRE-1a) on the positive strand and 5’-AGGCCA-3’ (TRE-2) located at the reverse strand of the HSV-1 TK promoter as the best half-sites for protein binding. The orientation of these half-sites for TRE-1 and TRE-2 was suggested as palindromic TREs with a six nucleotide in between (Pal 6). The point mutation changed the TRE-1a to 5’-AGGTGC-3’ and TRE-1b to 5’-GCGGCA-3’ on the reverse strand. This shift would allow TRE-1b’ and TRE-2 to form a direct repeat separated by three nucleotides (DR 3), presumably to generate positive regulation. B. PDBSum hydrogen bonding analysis of TRβ and RXRα hydrogen bonding to 5’-AGGTGA-3’ (wt TRE-1a), 5’-AGGTGC-3’ (mt TRE-1a’), and 5’-GCGGCA-3’ (mt TRE-1b’) suggested strongest binding to TRE-1b’ after the mutation due to one more hydrogen bond established from Arg160 of the RXRα.

**Fig. 2:**
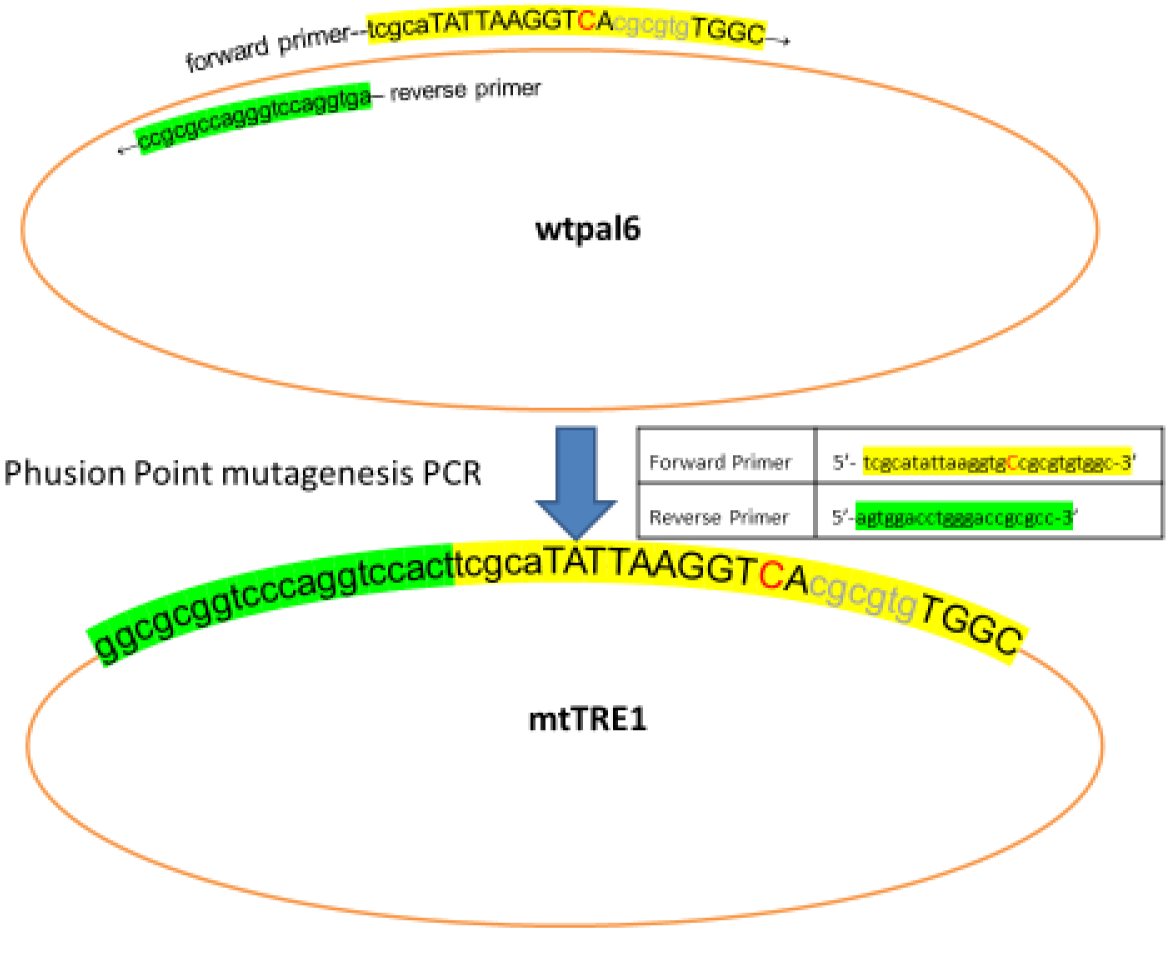
Site-directed mutagenesis. A. HSV-1 TK TREs and sequences of the mutant TREs. A 32-base of HSV-1 TK promoter regions from TATA box through the transcription initiation site was shown. The TATA box is framed and bases matching TRE half-site consensus sequences are identified by asterisks (*). TRE halfsite directions are identified by arrows. TREs are described and classified in reference to the well described TRE consensus sequence AGGT(G/C)A. The wt HSV-1 TK TRE is described as a palindrome with 6-nucleotide spacing (Pal6). B. Mutagenesis strategy. Specific primers were designed to introduce the mutation into the precise position using Phusion Site-Directed Mutagenesis system.

### Cells

The human prostate cancer cell line, LNCaP, was obtained from ATCC (Cat#: CRL-1740) and grown in RPMI-1640 supplemented with 10% FBS. Vero cells (ATCC Cat#: CCL-81) were grown in DMEM supplemented with 10% FBS. Cells were grown and maintained at 37 °C and 5% CO2 in a cell culture incubator.

### Neuroendocrine differentiation (NED)

The differentiation was achieved by removing androgen from the culture condition. In short, proliferating LNCaP cells were cultured in phenol red-free RPMI 1640 supplemented with 10% charcoal dextran treated FBS. The cells were seeded onto culture dishes at 4.0 × 10^3^ cells per cm^2^ of culture dish growth area. These conditions were maintained for at least 5 days before being treated further. A detailed protocol was reported previously (Figliozzi et al., 2014).

### Transfection

Lipofectamine 3000 (Cat#: L3000, Life Technologies) was used for transfection of LNCaP cells. Detailed protocol was provided by the manufacturer.

### Reporter assays

Luciferase activity was measured by a luminometer using the Dual-Luciferase reporter assay system (Cat#: PR-E1910, Promega). The cell lysate was collected for the luciferase assay after 48 h of transfection essentially described by the manufacturer. Luminescence was measured over a 10-s interval with a 2-s delay on the Synergy HTX multi-mode plate reader (Cat#: S1L, Biotek). The renilla luciferase activities were normalized against the internal firefly luciferase control driven by HSV-1 VP16 promoter to correct for transfection efficiency. The results were presented as the percent induction of the reporter plasmid in the presence or absence of T_3_ (100 nM).

### Antibodies

Detailed in Table 1. Anti-TRβ1 antibody was obtained from Thermo-Fisher (Cat#: MA1-215). Anti-RXRα (Cat#: ab41934) and anti-histone H3 tri-methylated K9 (Cat#: ab8898) was procured from Abcam Biotech. Anti-SP1 antibody was purchased from Santa Cruz Biotechnology (Cat#: SC-59).

**Table 1.**
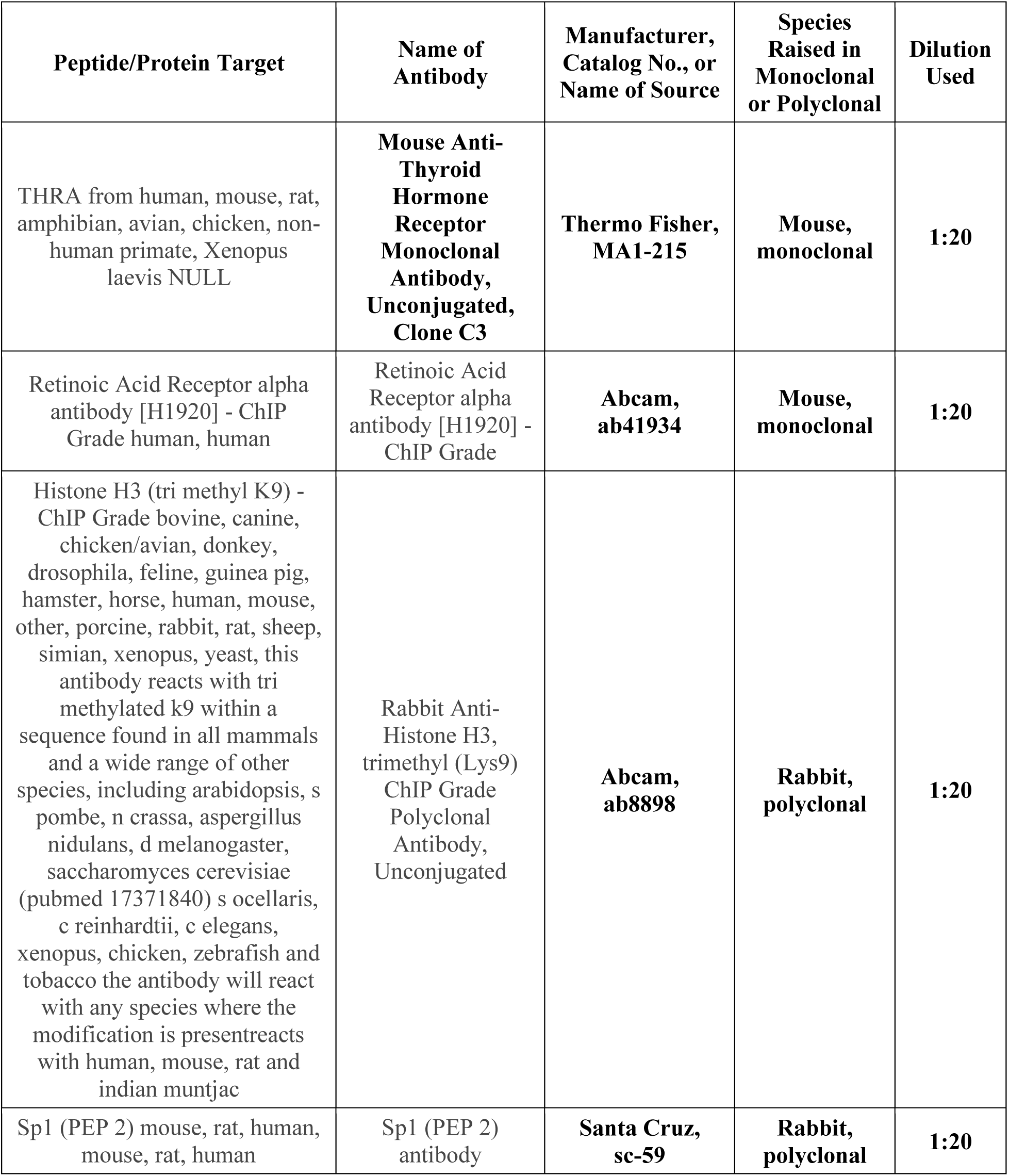
Antibodies.

| **Peptide/Protein Target** | **Name of Antibody** | **Manufacturer, Catalog No., or Name of Source** | **Species Raised in Monoclonal or Polyclonal** | **Dilution Used** |
| --- | --- | --- | --- | --- |
| THRA from human, mouse, rat, amphibian, avian, chicken, non-human primate, Xenopus laevis NULL | **Mouse Anti-Thyroid Hormone Receptor Monoclonal Antibody, Unconjugated, Clone C3** | **Thermo Fisher, MA1-215** | **Mouse, monoclonal** | **1:20** |
| Retinoic Acid Receptor alpha antibody [H1920] - ChIP Grade human, human | Retinoic Acid Receptor alpha antibody [H1920] - ChIP Grade | **Abcam, ab41934** | **Mouse, monoclonal** | **1:20** |
| Histone H3 (tri methyl K9) - ChIP Grade bovine, canine, chicken/avian, donkey, drosophila, feline, guinea pig, hamster, horse, human, mouse, other, porcine, rabbit, rat, sheep, simian, xenopus, yeast, this antibody reacts with tri methylated k9 within a sequence found in all mammals and a wide range of other species, including arabidopsis, s pombe, n crassa, aspergillus nidulans, d melanogaster, saccharomyces cerevisiae (pubmed 17371840) s ocellaris, c reinhardtii, c elegans, xenopus, chicken, zebrafish and tobacco the antibody will react with any species where the modification is present reacts with human, mouse, rat and indian muntjac | Rabbit Anti-Histone H3, trimethyl (Lys9) ChIP Grade Polyclonal Antibody, Unconjugated | **Abcam, ab8898** | **Rabbit, polyclonal** | **1:20** |
| Sp1 (PEP 2) mouse, rat, human, mouse, rat, human | Sp1 (PEP 2) antibody | **Santa Cruz, sc-59** | **Rabbit, polyclonal** | **1:20** |

### Electro-mobility Shift Assays (EMSA) and protein extraction

EMSA was performed using the LightShift™ Chemiluminescent EMSA Kit (Cat#20148, Fisher Scientific). The manufactures protocol was followed using 3’-biotin modified oligos corresponding to the wild type and TRE1 mutant sequences. The oligos were annealed and incubated with protein extracts from undifferentiated or differentiated LNCaP cells pretreated or without pretreatment of T_3_ in binding buffer for 30 min. The binding reaction mixtures were loaded onto 6% DNA Retardation Gel (Cat#: EC6365BOX, Invitrogen) and ran at 100V. The samples were transferred onto positively charged nylon membranes overnight using the capillary method. The membranes were blocked, washed, and exposed per the manufacturer. The membranes were sealed in blotting envelopes and photographed in a Bio-Rad ChemiDoc Imaging system. The sequences of the oligos are TK-Pal6-TRE-for: 5’-TAT TAA GGT GAC GCG TGT GGC CTC GAA CAC CG-3’ [BioTEG-Q] and TK-TRE1-mut1; 5’-TAT TAA GGT GCC GCG TGT GGC CTC GAA CAC CG-3’ [BioTEG-Q]. Their complementary sequences were synthesized and annealed using protocol described previously (Bedadala et al., 2010).

Protein Extracts were prepared from undifferentiated or differentiated LNCaP cells grown in T75s treated overnight with or without T_3_. Protein was isolated using RIPA buffer (Thermo Scientific cat# 89900) based on the protocol of the manufacturer. Briefly, the cells were washed twice with cold DPBS and exposed to cold RIPA buffer with Halt protease inhibitor cocktail (Thermo Sci cat# 78410) for 5 min on ice with gentle rocking. The lysate was scraped and collected followed by centrifuging at 14,000 × g for 15 minutes to pellet the cell debris. The supernatant was used for EMSA assays.

### Chromatin Immuno-precipitation (ChIP)

The ChIP was performed using ChromaFlash High-Sensitivity ChIP Kit from Epigentek, Farmingdale, NY (Cat#: P-2027-48). The protocol was described per the manufacturer. In short, test antibodies were first bound to Assay Strip Wells as well as Anti-RNA polymerase II (positive control) and non-immune IgG (negative control). The cells were subjected to cross-linking by adding media containing formaldehyde to a final concentration of 1% with incubation at room temperature (20–25 °C) for 10 min on a rocking platform (50–100 rpm). Glycine (1.25 M) was added to the cross-link solution (1:10) to stop the cross-linking. After appropriate mixing and ice-cold PBS washing and centrifuging, Working Lysis Buffer was added to re-suspend the cell pellet and incubate on ice for 10min. After carefully removing the supernatant, ChIP Buffer CB was added to re-suspend the chromatin pellet. Shear chromatin using Water Bath Sonication (EpiSonic 1100 Station, Cat No. EQC-1100, Epigentek). The program was set up at 20 cycles of shearing under cooling condition with 15 s On and 30 s Off, each at 170–190W. ChIP samples were centrifuged at 12,000 rpm at 4 °C for 10min after shearing and transferred supernatant to a new vial. Next set up the reactions by adding the ChIP samples to the wells that are bound with test antibodies, positive control, or negative control. The reaction incubation condition was 4°C overnight. ChIP samples then were washed per the protocol and subjected to reverse cross-linking at 60 °C for 45 min. Finally, the DNA samples were purified by spin column for quantitative PCR (qPCR).

### ChIP-qPCR

Quantitative analyses of ChIP and gene expression were performed by qPCR using myiQ SYBR green super-mix and iScript One-Step RT-PCR kits (BIO-RAD). Experiments were performed in triplicate with one set of primers per reaction. The ChIP primer sequences for TK TRE are 5′-ATG GCT TCG TAC CCC TGC CAT-3′ and 5′-GGT ATC GCG CGG CCG GGT A-3′. The qPCR reactions were carried out at 94 °C for 3 min, followed by 30 cycles of 94 °C for 15 s, 69 °C for 15 s, and 72 °C for 15 s. The results were calculated using Percent Input method with pre-immune antibody background subtracted.

## Results

### Computer analyses of putative binding of TR to alternative TRE

Swiss PDB Viewer (SPV) was used to generate two pdb files from 2NLL, each with either the DBD of TRβ or RXRα bound to its half-site. The generated pdb files were used as inputs for PiDNA. PiDNA randomly mutated the half-sites to yield hexamers with altered RXR or TR occupancy based on increased energy release due to binding and position frequency matrix analyses. PiDNA identified 5’-AGGTGA-3’ (TRE-1a) on the positive strand of the wt HSV-1 TK promoter (Fig. 1A) and 5’-AGGCCA-3’ (TRE-2) on the reverse strand as the optimal half-sites for protein binding (data not shown). The orientation of these half-sites suggests a palindromic TRE with a six-base pair spacer (Pal 6) as suggested by us and others (Figliozzi et al., 2014;Hsia et al., 2010;Maia et al., 1996;Park et al., 1993). Interestingly, PiDNA also suggested an alternative binding to 5’-GCGTCA-3’ (TRE-1b) located at the reverse strand (Fig. 1A). This half-site exhibited the same binding strength in comparison to TRE-1a (data not shown).

Computational observations revealed that a TRE-1a 6A/C mutation on the positive strand to become 5’-AAGTGC-3’ would create a new TRE-1b’ with half-site sequence of 5’-GCGGCA-3’ on the reverse strand identified by PiDNA as preferentially bound by RXR in comparison to TRE-1a’ (Fig. 1B) (Rastinejad et al., 2013;Rastinejad et al., 1995). Thus, TRE-1b’ and TRE-2 would create a pair of direct TRE repeat with three nucleotide spacing, likely to behave as positive TRE conferring positive regulation. PDBSum analysis of TRβ and RXRα hydrogen bonding further suggests the likelihood of a direct repeat on the mutant promoter. The mutant TRE1b’ would gain one more hydrogen bond from Arg160 from the RXRα (Fig. 1B) to strengthen the interaction which is absent from the wild type.

### Site-directed mutagenesis

HSV-1 TK promoter contains a pair of TREs exhibiting palindrome repeats with six nucleotides separating each other (Pal6 TREs). These Pal6 TREs reside between the TATA box (47933–47937) and the transcription initiation site (47911) (Fig. 2) based on the HSV-1 complete genome sequence accession number X14112. This context was discussed previously to produce negative regulation by T_3_ in neural cells but different cells generating different opposite results even in several cells with neural origin (Figliozzi et al., 2014;Hsia et al., 2010;Maia et al., 1996;Park et al., 1993). To address the importance of TRE sequence in the T_3_-mediated negative regulation, a point mutation was introduced followed by reporter assays to study the regulatory effects. For easy discussion, the TRE adjacent to TATA box was named TRE1 and the other one was called TRE2 (Fig. 2). The mutation was introduced to the end of TRE1 from 5’-AGGTGA-3’ to 5’-AGGTGC-3’ (Fig. 2, mtTRE1). The original TK TRE was named wtPal6. The resulting plasmid mtTRE1 was confirmed by sequencing (data not shown).

### TK promoter and its mutant were not regulated by T_3_ in undifferentiated LNCaP cells

These reporter plasmids were first tested by dual luciferase (DLuc) assay after transfection of undifferentiated LNCaP cells treated with and without T_3_. DLuc assays showed that the T_3_ treatment did not affect the promoter activity of any plasmid (Fig. 3A), indicating that there is no T_3_-mediated regulation in undifferentiated cellular environment.

**Fig. 3:**
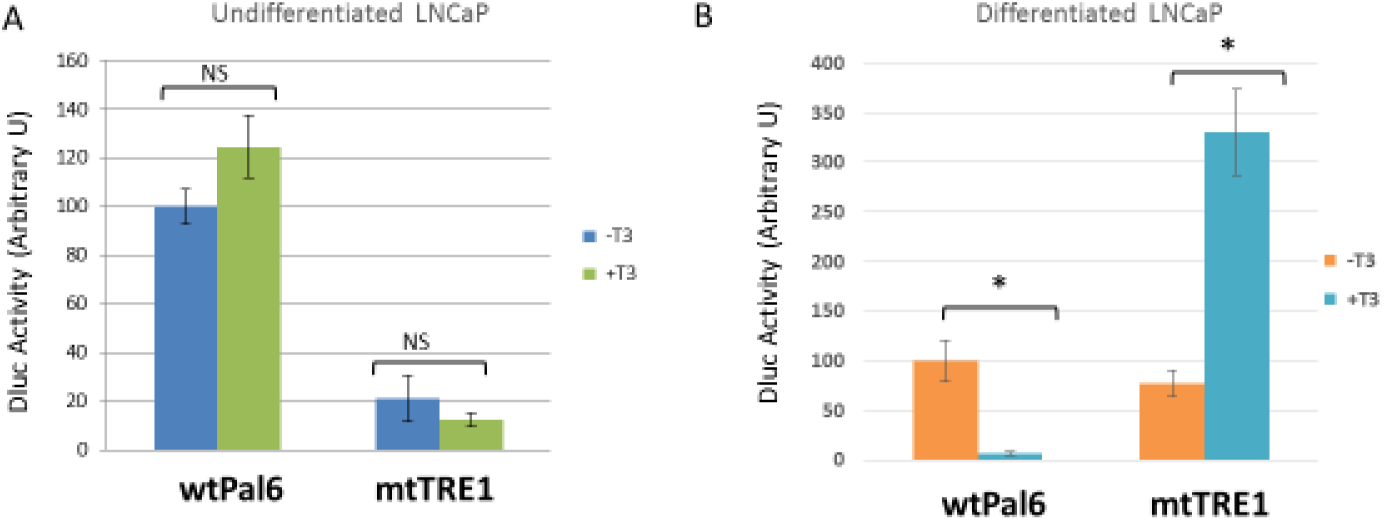
Transfection and DLuc assays. A. Transfection of undifferentiated cells. Dual-Luciferase assays of undifferentiated LNCaP cells transfected with HSV-1 TK TRE luciferase reporter plasmids: wtPal6 and mtTRE treated with and without T_3_, were performed in triplicate. The effects of T_3_ on promoter activity were not significant and wt promoter generally had stronger activity in comparison to mt counterpart. Two-way ANOVA with Holm-Sidak post-hoc analysis suggests that the differences in activity is not significant. B. Transfection of differentiated cells. Similar Dual-Luciferase assays of differentiated LNCaP cells transfected with the reporter plasmids in the presence or absence of T_3_, were done in triplicate. It appeared that mt plasmids exhibited different T_3_-mediated regulation compared to the wt version. Two-way ANOVA with Holm-Sidak posthoc analysis suggests that the differences in activity is statistically significant where * denotes p<0.05.

### TK single nucleotide mutant reporter plasmids exhibited distinct T_3_-mediated regulation in differentiated LNCaP cells

The plasmids were later transfected into differentiated LNCaP cells with or without T_3_ treatment. Fig. 3B shows T_3_ treatment caused a 10-fold reduction in wtPal6 promoter activity, consistent with our previous finding that T_3_ mediated a negative regulation in differentiated cellular background. In contrast mtTRE1 activity showed the opposite regulatory profile with a 4-fold increase upon T_3_ treatment (Fig. 3B, mtTREl panel). In differentiated LNCaP, under T_3_ the wt promoter activity was at least 70-fold weaker than mtTRE1 (Fig. 3B). Together these observations demonstrated that single nucleotide changes within the TREs can disrupt the normal T_3_-mediated repression of HSV-1 TK promoter only in differentiated cells.

### Addition of T_3_ to differentiated cells without prior treatment of T_3_ reduced its capacity of TR to bind wt TREs

The binding of TR to the TREs under the influence of hormone was investigated by EMSA. Wild type promoter oligo and protein extract from differentiated cells not pretreated with T_3_ were first tested. Shifted bands were detected where the addition of T_3_ slightly reduced the band intensity (compare Fig. 4A, lane 2 and 3). The decreased interaction by T_3_ was more significant if longer electrophoresis was allowed (Fig. 4C, lane 2 and 3). These observations demonstrated that proteins from the extract interacted with the TRE oligo and T_3_ reduced the binding. To address if TR contributed to the binding to the TRE, antibodies against TR or RXR were introduced in the EMSA. The results showed that the retardation bands were completely abolished upon the addition of anti-TR Ab and only slightly affected upon the addition of anti-RXR Ab (Fig. 4A, lane 4–5 and 6–7, respectively), supporting that TR played a critical role in this interaction, perhaps by direct binding to the TK-TREs with RXR functioning as a partner.

**Fig. 4:**
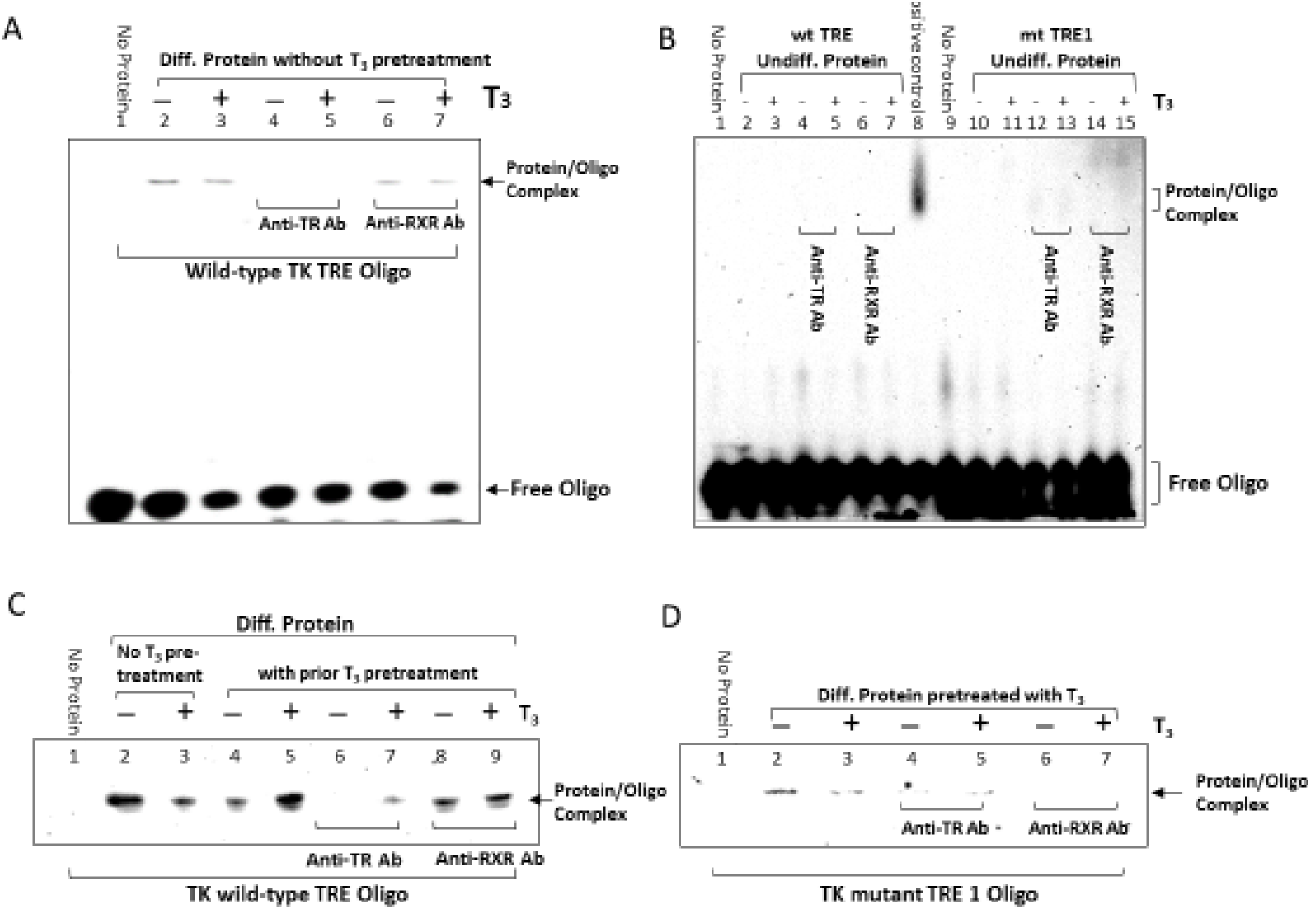
EMSA. A. The labeled wt TRE oligo was incubated with differentiated protein extract without hormone pretreatment followed by gel electrophoresis and the image was captured. Lane 1, no protein; Lane 2–7, incubated with protein extract from differentiated cells; Lane 4–5, with anti-TRβ Ab; Lane 6–7, with anti-RXRα Ab. The image came from a 10-well 6% DNA Retardation Gel. B. Undifferentiated protein extract was used for comparison. It is shown that TRE-TR interaction was not seen in undifferentiated extract. Both wtTRE (lane 2–7) and mtTRE1 (lane 10-15) failed to interact with TR. positive control was wtTRE interacted with differentiated protein extract without T_3_ pretreatment, like Fig. 4A, lane 2. Note that this gel had 15-well so the bands were not as sharp as the other gels, which were 10-well gels. C. The comparison of TREs oligo/TR interaction with or without T_3_ pretreatment was analyzed. Lane 2-3, extract without T_3_ pretreatment. Lane 4–7, extract with prior T_3_ pretreatment. It appeared that TRE/TR interaction increased in the presence of T_3_ with hormone pretreatment (compare lane 4 and 5). Anti-TR Ab failed to completely abolish the interaction (lane 6 and 7), suggesting a robust binding of TR to TRE in T_3_. The free oligos were run out of the gel due to increased electrophoresis time to improve the resolution. D. The interaction of mutant TRE1 to TR with pretreatment was investigated. Lane 1, no extract; Lane 2 and 4, no T_3_; Lane 3 and 5, with T_3_; Lane 4–5, anti-TR Ab added. The intensity of mtTRE1 was reduced in the presence of T_3_ (compare lane 2 and 3). Anti-TR Ab addition did not alter the interaction (lanes 4 and 5). Anti-RXR Ab addition altered the interaction (lanes 6 and 7). The free oligos were not seen because of elongated electrophoresis.

### TK-TREs were not bound by TR from undifferentiated cells

Protein extract from undifferentiated LNCaP cells of equivalent total protein concentrations used in differentiated extract experiments in Fig. 4A were used to compare the TRE/TR interaction. Surprisingly the strong interaction between wtTRE and TR from differentiated cells (Fig. 4B, lane 8) was not observed (Fig. 4B, lane 1–7) even though TR is detectable at this dilution (data not shown). The mutant TRE oligo yielded similar results of no interaction (Fig. 4B, lane 9–15). Previous study indicated that TRβ was expressed in undifferentiated LNCaP cells (Figliozzi et al., 2014). This results suggested that differentiated status played critical roles in TR binding to TK TREs.

### Interaction of T_3_ to TR of differentiated cells pretreated with T_3_ increased its binding capacity to wt TREs

To mimic the physiological condition, we pretreated differentiated LNCaP cells with T_3_ for 48 hours and wash the cells with PBS followed by protein extraction (prior T_3_ pretreatment). It was noted that T_3_-pretreated protein extracts caused the band intensity to increase upon T_3_ addition (compare Fig. 4C, lane 4 and 5). Upon addition of anti-TR Ab, the lane lacking T_3_ binding treatment nearly disappeared (Fig. 4C, lane 6) whereas its T_3_ treated counterpart band was still clearly visible with reduced intensity (Fig. 4C, lane 7), suggesting that ligand TR, if pretreated with hormone, exhibited a stronger binding capacity to TREs. The addition of anti-RXR Ab had little effect on these bands (Fig. 4C, Lane 8 and 9).

### mtTRE1 demonstrated different pattern of binding in comparison to wt TREs

TRE mutants were investigated for their binding by TR. It was shown that mtTRE1 band intensity decreased to very low level in the presence of T_3_ (compare Fig. 4D, lane 2 and 3), similar to the pattern of no prior T_3_ treatment (compare Fig. 4A, lane 2 and 3). Anti-TR Ab experiments again suggested that this interaction required TR (Fig. 4D, lane 4 and 5). Unlike the wtTRE, the addition of anti-RXR Ab also reduced the band signal, which suggested that RXR was also involved in DNA binding on the mtTRE (Fig. 4C, Lane 6 and 7). Together these observations suggested that the mutation disrupted the interaction of nuclear hormone receptors to the wild-type TRE thus altering the negative regulation.

### Repressive chromatin H3K9me3 enriched at wt promoter in the presence of T_3_ was released in mtTRE1 promoter

ChIP was performed to investigate the impact of single nucleotide change of the TREs on the chromatin recruitment to the promoter. In this experiment using an antibody against H3K9me3, a repressive histone previously reported to be associated with TK promoter with T_3_ (Figliozzi et al., 2014), we showed that in the wt promoter the tri-methylated histone interaction was increased 4.3-fold upon T_3_ addition (Fig. 5A), in agreement with our hypothesis that T_3_ mediated repression in differentiated cells. In contrast the opposite effect was shown for the mtTRE1, where H3K9me3 recruitment decreased by as many as 5.5-fold in the presence of T_3_ (Fig. 5A). This result indicated that one nucleotide mutation shifted the histone profile under the same condition.

**Fig. 5.**
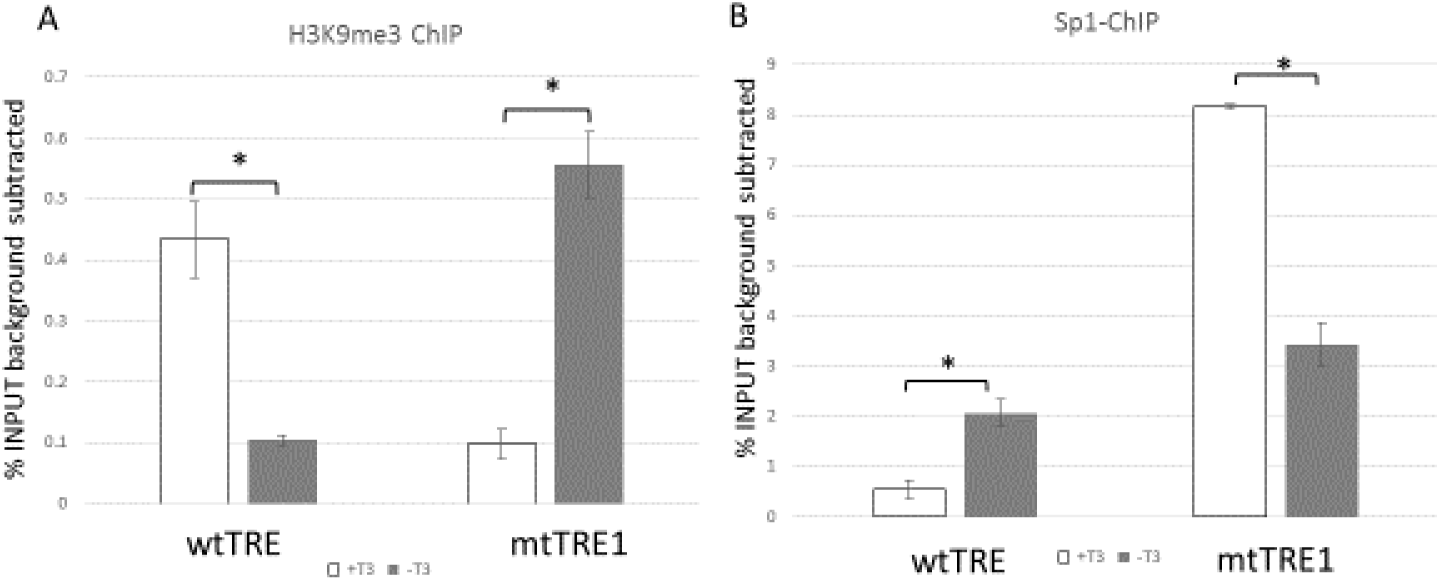
a: Enrichment of repressive chromatin H3K9me3 at the HSV-1 TK TREs. ChIP assay measuring the enrichment of H3K9me3, a repressive chromatin protein marker, to HSV-1 TK TREs from differentiated LNCaP cells transfected with wtTRE and mtTRE1 treated with and without T_3_ presented as presented as percent input with IgG background subtracted. T_3_ treatment of wtTRE resulted in a 4.5-fold increase in comparison to no T_3_ treatment. The mtTRE1, however, showed an opposite effect where T_3_ treatment caused a reduction. **b: Recruitment of Sp1 to the TK TREs** The occupancy of transcription factor Sp1 to HSV-1 TK TREs under the influence of T_3_ was analyzed by ChIP from differentiated LNCaP cells transfected with wtTRE and mtTRE1. The results were presented as percent input with IgG background subtracted. For the wtTREs the T_3_ treatment resulted in a 4-fold reduction of Sp1 recruitment comparing to no T_3_ treatment. The mtTRE1 showed a reversed effect that T_3_ increased the Sp1 occupancy in comparison to no T_3_.

### Transcription factor Sp1 occupancy was reversed by TRE sequence changes

Recruitment of transcription factor was studied by ChIP Assays using antibodies against Sp1. It was revealed that Sp1 recruitment to the wt promoter increased 4-fold without T_3_, whereas decreased approximately 3-fold in the mtTRE1 promoter (Fig. 5B). There are two Sp1 binding sites, -111 and −59 bp, respectively (Coen et al., 1986) and they did not overlap with TK TREs. This observation suggested that mutant TREs may alter chromatin configuration therefore influence the interaction of transcription factor to its binding sites.

## Discussion

Seemingly minute TRE sequence differences in promoters would direct chromatin recruitment and determine the type of T_3_-mediated regulation. However, the mechanisms are poorly understood. Putative TREs were identified within a wide variety of promoters and characterized as *bona fide* TREs by molecular analyses. Several TREs were depicted in viral promoters including HSV-1 TK (Bedadala et al., 2010;Desai-Yajnik and Samuels, 1993;Desvergne and Favez, 1997;Hsia et al., 2011;Hsia and Shi, 2002;Hsia et al., 2003;Hsia et al., 2001;Zuo et al., 1997). For decades the HSV-1 TK promoter has been used as a transcriptional control and studied for its T_3_ sensitivity (Maia et al., 1996). Bioinformatics demonstrated a pair of nontraditional palindromic TK TREs located between TATA box and the transcription initiation site, in a context like the thyroid stimulating hormone alpha (TSHα) TREs, one of the most well characterized negative TREs (Carr et al., 1992;Hollenberg et al., 1995;Jacobs and Kuhn, 1992;Kohn et al., 1992;Rentoumis et al., 1990). It is thus hypothesized that TK TREs would be bound by TR/T_3_ to confer negative regulation in differentiated cells with neural phenotype but not non-neural cells (Hsia et al., 2010;Park et al., 1993). Nonetheless it is quite complicated. First it was shown that TR/T_3_ exhibited no regulation of TK transcription on most of the non-neural cells (Hsia et al., 2011;Maia et al., 1996) but generated good negative regulation in differentiated mouse neuroblastoma cells N2a (Hsia et al., 2010). These observations supported the hypotheses nicely. However, more studies indicated that there was no T_3_-mediated regulation in rat pituitary GH4C1 cells (Park et al., 1993), a popular model to investigate molecular mechanisms of TRH receptor function, signal transduction, electrophysiological studies on plasma membrane calcium channels, and intracellular calcium homeostasis in pituitary cells. Nevertheless, it appeared that TR/T_3_ delivered negative regulation in human choriocarcinoma cell line JEG-3, a placental epithelial-trophoblast-like cells had little evidence of differentiation (Maia et al., 1996). Furthermore, HSV-1 TK was negatively regulated by T_3_ in a differentiated but not a true neural cells LNCaP cells (Figliozzi et al., 2014). Together these statements suggested that T_3_-mediated regulation on a promoter with palindromic TREs may be controlled by multiple factors such as cell origin, host regulatory protein recruitment, TRE sequence composition/context, nuclear receptor subtypes, dimerization preferences, differentiation status, and chromatin context.

The negative regulation of T_3_ and TR is not as well-characterized as the positive regulation but is of equal importance. A number of hypotheses have been suggested such as suppression of Sp1 stimulation (Xu et al., 1993), cAMP response element binding (CREB) competition (Mendez-Pertuz et al., 2003), recruitment of chromatin insulator protein CTCF (Burke et al., 2002), the downstream binding to TATA box and direct interaction of TFIID (Crone et al., 1990), hetero-to homodimer conformational alteration (Bendik and Pfahl, 1995), interaction of GATA2 and TRAP220 dissociation (Matsushita et al., 2007), TR/Sp1 competition in the first exon (Villa et al., 2004), ligand-mediated recruitment of histone deacetylases (HDAC) complex (Sasaki et al., 1999), interaction of TR/TATA binding protein (TBP) and HDAC (Sanchez-Pacheco and Aranda, 2003), and conversion of corepressor SMRT to coactivator by TR (Berghagen et al., 2002). It is being debated that if T_3_-mediated negative regulation required TR-DNA binding (Madison et al., 1993;Shibusawa et al., 2003;Tagami et al., 1999). In this study, another layer of complication, differentiation, was added to the sophisticated regulation. It is known that T_3_ induces differentiation in several different tissues of many different species (Nygard et al., 2003). Studies suggested that T_3_ induced withdrawal from the cell cycle through S-phase genes inhibition such as E2F1, S-phase-specific DNA polymerase alpha, thymidine kinase, and dihydropholate reductase genes. For instance, ligand TR triggered differentiation by suppressing E2F-1 gene expression, a key transcription factor that controls G1-to S-phase transition (Nygard et al., 2003). It is surprising to learn that differentiated cells with prior treatment of T_3_ exhibited stronger TR binding capacity to wtTRE in the presence of T_3_. The exact mechanisms are not understood. It is likely that prior exposure to T_3_ changed the gene expression profiles to modulate the TR/TRE interaction. It cannot be ruled out that differentiated and undifferentiated conditions generated distinct chromatin environments allowing these discrete interactions.

The occupancy of transcription factor Sp1 to the TK promoters with different TREs was assessed. The ChIP results matched the Dluc assays and indicated that Sp1 occupancy promoted active transcription in the absence of T_3_ (compare Fig. 3B and Fig. 5B). It is worth noting that in the presence of T_3_, Sp1 recruitment to mtTRE1was enhanced by 16-fold in comparison to wtTRE (Fig. 5B). It seemed that Sp1 binding was determined, at least in part, by TR binding (compare Fig. 5B to Fig. 4C and 3D) and the ligand appeared to play a critical role. Position weight matrix comparison of the wild type and mutant promoter for Sp1 binding sites did not reveal additional sites upon mutation (data not shown). Two Sp1 binding sites are not found overlapping the TK TRE but well upstream of the TATA box (-111 bp and −59 bp), therefore the mutations did not affect the Sp1 binding site but still altered its binding profile (Coen et al., 1986). It is unclear how the mutation achieved this phenomenon. We therefore speculate that in both cases T_3_ modulated histone modification and the resulting chromatin environment at this promoter determined Sp1 occupancy.

HSV-1 TK is not essential during lytic infection (Coen et al., 1986;Knipe and Howley, 2013) however it plays a critical role in the drug action of acyclovir and reactivation (Kosz-Vnenchak et al., 1993;Nichol et al., 1996;Tal-Singer et al., 1997;Valyi-Nagy et al., 1994;Wilcox et al., 1992). In resting cells such as neurons dNTPs are absent and requires TK action to provide dNTPs for viral replication (Knipe and Howley, 2013). In contrast to the lytic infection that viral replication is not essential for α and β gene expressions, TK was suggested to promote replication followed by efficient α and β gene expressions in neurons during reactivation to complete the life cycle (Nichol et al., 1996;Tal-Singer et al., 1994). In addition, TK null mutant showed decreased viral gene expression upon reactivation (Kosz-Vnenchak et al., 1993) and in vivo reactivation studies revealed that TK was among the first genes to be expressed (Pesola et al., 2005;Tal-Singer et al., 1994).

In conclusion, the unique negative TREs of HSV-1 TK were characterized in this report by side-directed mutagenesis. T_3_ was not sufficient to control TK promoter activity in undifferentiated condition but conferred repression while cells were differentiated. A point mutation at TRE1 close to TATA box changed the T_3_-mediated negative regulation into positive fashion. In vitro EMSA suggested this wide spaced TREs would favor TR monomer binding and strong TR interaction to the TREs particularly in the presence of T_3_ during differentiation. Ligand TR under differentiation appeared to suppress TK transcription by creating a repressive chromatin environment and by reducing the recruitment of the important transcription factor Sp 1 to the promoter. This understanding of this complex regulation may have implication to control alpha herpes virus infection and reactivation.

## Abbreviation

HSV-1: Herpes Simplex Virus Type-1
T_3_: Thyroid Hormone
qPCR: quantitative Polymerase Chain Reaction
TRβ1: Thyroid Hormone Receptor Beta 1
TRE: Thyroid Hormone Receptor Element
EMSA: Electro Mobility Shift Assay
ChIP: Chromatin Immuno-Precipitation
Sp1: specificity protein 1
RXR: Retinoid X Receptor
H3K9me3: histone H3 lysine 9 tri-methylated
VZV: Varicella Zoster Virus
TK: thymidine kinase
DLuc: Dual Luciferase

## Acknowledgement

This project is supported by R01NS081109 to SVH from NIH and the content is solely the responsibility of the authors and does not necessarily represent the official views of the NINDS/NIH. The authors appreciate the assistance of the editorial staff at UMES.

## Author contributions

RWF and SVH initiated this investigation and designed the study. FC provided the cultured cells. RWF did all transfection experiments and analyzed the promoter activity. RWF performed the EMSA. FC designed the primers and performed all the ChIP analyses. RWF performed the computational approach. SVH wrote the original draft of the manuscript. RWF and FC contributed to portion of the manuscript writing. All authors analyzed and validated the results and approved the final version of the manuscript.

